# Deep Learning-Enabled Point-of-Care Sensing Using Multiplexed Paper-Based Sensors

**DOI:** 10.1101/667436

**Authors:** Zachary Ballard, Hyou-Arm Joung, Artem Goncharov, Jesse Liang, Karina Nugroho, Dino Di Carlo, Omai B. Garner, Aydogan Ozcan

## Abstract

We present a deep learning-based framework to design and quantify point-of-care sensors. As its proof-of-concept and use-case, we demonstrated a low-cost and rapid paper-based vertical flow assay (VFA) for high sensitivity C-Reactive Protein (hsCRP) testing, a common medical test used for quantifying the degree of inflammation in patients at risk of cardio-vascular disease (CVD). A machine learning-based sensor design framework was developed for two key tasks: (1) to determine an optimal configuration of immunoreaction spots and conditions, spatially-multiplexed on a paper-based sensing membrane, and (2) to accurately infer the target analyte concentration based on the signals of the optimal VFA configuration. Using a custom-designed mobile-phone based VFA reader, a clinical study was performed with 85 human serum samples to characterize the quantification accuracy around the clinically defined cutoffs for CVD risk stratification. Results from blindly-tested VFAs indicate a competitive coefficient of variation of 11.2% with a linearity of R^2^ = 0.95; in addition to the success in the high-sensitivity CRP range (i.e., 0-10 mg/L), our results further demonstrate a mitigation of the hook-effect at higher CRP concentrations due to the incorporation of antigen capture spots within the multiplexed sensing membrane of the VFA. This paper-based computational VFA that is powered by deep learning could expand access to CVD health screening, and the presented machine learning-enabled sensing framework can be broadly used to design cost-effective and mobile sensors for various point-of-care diagnostics applications.

## INTRODUCTION

Computation has great potential for improving diagnostics. By identifying complex and nonlinear patterns from noisy inputs, computational tools present an opportunity for automated and robust inference of medical data. For example, several studies have shown deep learning as a method to automatically identify tumors from an image, potentially enabling diagnostics in low-resource settings that lack a trained diagnostician ^1–3^. Additionally, computational solutions have been demonstrated earlier in the diagnostics pipeline to virtually stain pathology slides and enhance image resolution through the use of convolutional neural networks ^4–6^. Though much of this recent success is within the field of imaging, diagnostics that rely on biosensing can similarly leverage computational tools to improve sensing results and design future systems.

Point-of-care (POC) testing can especially benefit from computational sensing approaches. Due to their low-cost materials, compact designs, and requirement for rapid and user-friendly operation, POC tests are often less accurate when compared to traditional laboratory tests and assays ^7–12^. For example, paper-based immuno-assays such as rapid diagnostic tests (RDTs) offer an affordable and user-friendly class of POC tests which have been developed for malaria, HIV-1/2, and cancer screening, among other uses ^13–17^. However, these RDTs lack the sensitivity and specificity needed for certain diagnostic applications largely due to issues of reagent stability, fabrication and operational variability, as well as matrix effects present in complex samples such as blood ^15,16,18^. Additionally, a well-known competitive binding phenomenon called the hook-effect can lead to false reporting of results, specifically in instances where the sensing analyte can be present over a large dynamic range ^19–24^. Therefore, computational tools alongside portable and cost-effective assay readers present a unique opportunity to compensate for some of these constraints ^25–31^. By quantifying the signals generated on paper-based substrates, machine learning algorithms have the potential to significantly improve the performance of POC sensors, without a significant hardware cost or increased complexity to the assay protocol.

As a unique demonstration of this emerging opportunity at the intersection of computational sensing and machine learning, here we report a computational paper-based vertical flow assay (VFA) for cost-effective high-sensitivity C-Reactive Protein (hsCRP) testing, also referred to as cardiac CRP testing (cCRP) ^32^. This low-cost and rapid (< 12 min) VFA uses a multiplexed sensing membrane and diagnostic algorithm based on neural networks to accurately quantify CRP concentration in the high-sensitivity range (i.e. 0-10 mg/L), as well as to identify samples outside of this range despite the presence of the hook-effect.

CRP is a general biomarker of inflammation, however slightly elevated CRP levels in blood can be an indicator of atherosclerosis, and have been shown to be a predictor for heart attacks, stroke, and sudden cardiac death for patients with and without a history of CVD ^33–36^. Therefore, the hsCRP test is a quantitative test commonly ordered by cardiologists to stratify certain patients into low, intermediate, and high risk groups for CVD based off of clinically defined cut-offs: below 1 mg/L is considered low risk, between 1 and 3 mg/L is intermediate risk, and above 3 mg/L is high-risk ^37^. As a result, the hsCRP test requires a high degree of accuracy and precision, especially around the clinical cut offs, putting it out-of-reach of traditional paper-based systems ^38^. Additionally, in the presence of infection, tissue injury, or other acute inflammatory events, CRP levels can rise nearly three orders of magnitude, making hsCRP testing with immuno- and nephelometric-assays vulnerable to the hook-effect ^34,36,39^. As a result, samples with greatly elevated CRP levels can be falsely reported as within the hsCRP range (i.e., < 10 mg/mL), and therefore wrongly interpreted for CVD risk stratification.

To address these existing challenges of POC hsCRP testing, in this work we implemented a computational VFA-based sensing framework to jointly develop the CRP quantification algorithm and multiplexed sensing membrane configuration, computationally selecting the most robust subset of sensing channels with which we accurately infer the CRP concentration. We performed a clinical study with 85 patient serum samples and >250 VFA tests created over multiple fabrication batches, and compared the sensor performance to an FDA-approved assay and nephelometric reader (Dimension Vista System, Siemens). Our blind testing results yielded an average coefficient of variation (CV) of 11.2% and a coefficient of determination (R^2^) of 0.95 over an analytical measurement range of 0 mg/L to 10 mg/L. It is important to note that although there is no FDA-approved POC hsCRP sensor, various systems have been demonstrated in the literature^40–44^. However, the tests which report accurate quantification in the high-sensitivity range employ fluorescent-based chemical assays and benchtop readers to overcome the performance limits of their traditional colorimetric counterparts. In contrast, this work uniquely demonstrates a new data-driven sensor design and read-out framework, powered by deep learning, for improving POC testing. We applied this machine learning-enabled sensing framework to a colorimetric paper-based multiplexed test for quantification of hsCRP as a use-case, and demonstrated its competitive quantitative performance using a mobile reader, without the need for more advanced and sensitive molecular assays and their corresponding benchtop read-out systems.

We believe that the presented POC hsCRP sensor platform could provide a rapid and cost-effective means to obtain valuable diagnostic and prognostic information for CVD, expanding access to actionable health information, especially for at-risk populations that often go underserved (34,35). Broadly, our results also highlight computational sensing as an emerging opportunity for iterative assay and sensor development. Given a training data set, machine learning-based feature selection algorithms can be implemented to determine the most robust sensing channels for a given multiplexed system such as protein micro-array, well-plate assay, or multi-channel fluidic device, among others. This can therefore lead to optimized and cost-effective implementations of multiplexed bio-sensing systems for future POC diagnostic applications.

## METHODS

### Multiplexed VFA

#### Overview

The multiplexed VFA platform is comprised of functional paper layers stacked within a 3D-printed plastic cassette. These layers contain different paper materials and wax printed structures which have been optimized to support uniform vertical flow of serum across a two-dimensional nitrocellulose sensing membrane (Fig. 1a, Table S1). Similar to conventional paper-based immunoassays, the VFA works by immobilizing a target analyte onto a paper substrate through binding to a complimentary capture antigen or antibody previously adsorbed within the porous structure ^45^. Gold nanoparticles conjugated with a secondary antibody are then introduced and bound to the immobilized analyte in a sandwich structure, resulting in a color signal on the sensing membrane. The operation of our VFA test involves three sequential injection steps: 1) the running buffer, 2) the sample serum and nanoparticle conjugate, and 3) the washing buffer (Fig 1c). After a 10 minute wait-period, the assay is complete and the VFA cassette is opened by twisting apart the top and bottom case, revealing the multiplexed sensing membrane on the top layer of the bottom case (Fig. 1c). This bottom case is then inserted into a custom-designed mobile-phone reader. An image of the activated multiplexed sensing membrane is subsequently captured and analyzed via a fully-automated image processing and deep learning-based CRP quantification algorithm (Fig. 1d).

**Figure 1.**
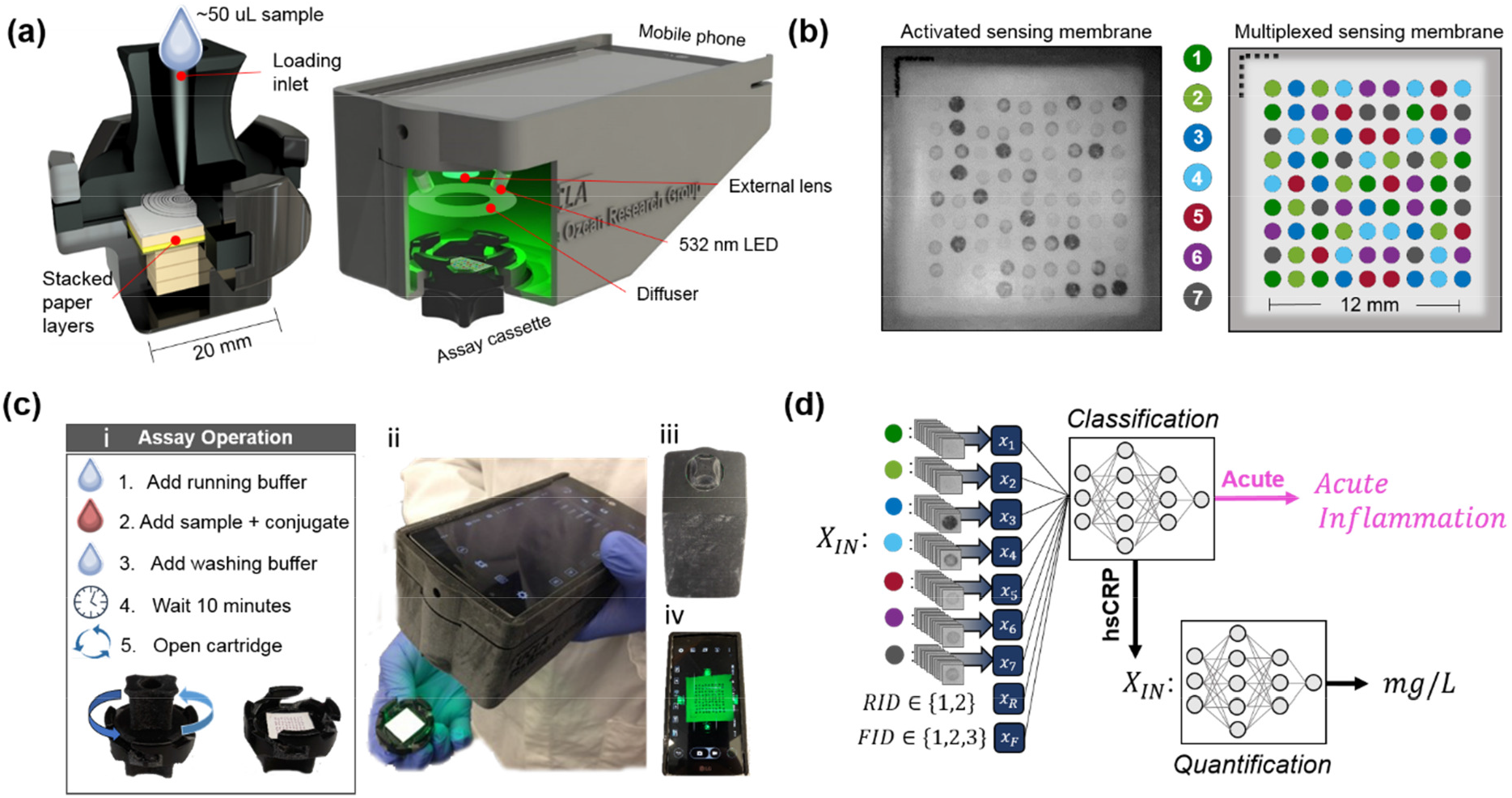
(a) The multiplexed vertical flow assay (VFA) cassette cross-section and mobile-phone reader with the inserted VFA cassette to be tested. (b) The multiplexed sensing membrane contained within the VFA cassette. The algorithmically determined immunoreaction spot layout (right) contains seven unique spotting conditions, each of which uniquely reacts with the sensed analyte and the signal-forming Au NPs. A raw image of an activated sensing membrane taken with the mobile-phone reader is shown to the left. (c) (i) The VFA assay operation protocol (ii) The VFA cassette and mobile phone reader after the assay completion. The VFA cassette inserted into the mobile phone reader from the (iii) bottom and (iv) top view. (d) Block diagram of the computational analysis, showing the input features *X_IN_* which contain, the average signals from like-spotting conditions along with the reagent batch ID (RID) and the fabrication batch ID (FID).

#### Multiplexed sensing membrane fabrication and VFA assembly

The multiplexed sensing membrane contains up to 81 spatially isolated immunoreaction spots that are each defined by a ‘spotting condition’ which refers to the capture protein and the associated buffer dispensed onto the nitrocellulose sensing membrane prior to assembly and activation. Therefore, to design the multiplexed sensing membrane for computational analysis, a custom spot-assignment algorithm was developed to generate a ‘spot map’ within the active area of the sensor. Based on a given grid spacing and number of spotting conditions, the assignment algorithm distributes spotting conditions such that no single spotting condition is disproportionately positioned near the center or the edge of the sensing membrane. Because the vertical flow rate can vary radially across the sensing membrane, leading to variations of each immunoreaction across the sensor area, this step mitigates a potential bias on any given spotting condition. With seven spotting conditions (see Table S2.) in a 9×9 grid format (1.3 mm periodicity), the spot-assignment algorithm produced the map shown in Figure 1c, which was implemented as the initial design for this study.

An automated liquid dispenser (MANTIS, Formulatrix^®^) was used to deposit 0.1 μL of the different protein conditions directly onto a nitrocellulose (NC) membrane in the algorithmically determined pattern shown in Fig. 1c. During the spotting process, up to 24 NC sensing membranes were produced on a single connected sheet, constituting one fabrication batch, and up to three batches were produced on a given day. In order to evaluate batch-to-batch variations, we intentionally produced sensing membranes over multiple fabrication batches as well as with two reagent batches (*i.e*. sets of reagents which had unique storage times and/or lot numbers). Each sensing membrane was therefore tagged with a corresponding fabrication batch ID (FID, *e.g*., 1, 2 or 3,) and reagent batch ID (RID, *e.g*., 1 or 2).

Following the automated spotting procedure, the NC sheets were incubated at room temperature for 4 hours after which they were submerged in 1% BSA blocking solution and allowed to incubate at room temperature for 30 min. The NC sheets were then dried in an oven at 37°C for 10 min, after which they were cut into individual sensing membranes (1.2 x 1.2 cm) using a razor. The remaining paper materials contained in the VFA were produced following the methods outlined in a previous publication ^45^. All the paper materials, including the NC sensing membrane were then assembled within the top and bottom cases of a 3-D printed VFA cassette, with foam tape holding together the paper stack (see Table S1).

#### hsCRP assay procedures

Each hsCRP measurement with our VFA test is performed as follows: first 5 μL of serum sample is diluted 10 times in a running buffer (3% tween 20, 1.6% BSA in PBS) resulting in a 50 μL sample solution. Then 200 μL of running buffer is injected into the VFA inlet and allowed to absorb. After absorption into the VFA paper-stack (~ 30 sec), 50 μL of sample solution is mixed with 50 μL of the gold-nanoparticle (Au NP) conjugate solution (see Supplementary Information for synthesis), and the mixture is pipetted into the inlet and allowed to absorb. Lastly, after absorption of the sample solution, 400 μL of the running buffer is added to wash away the nonspecifically bound proteins and Au NPs. After a 10 minute reaction time, the VFA cassette is then opened, and inserted into the bottom of the mobile-phone reader (Fig 1a.). This mobile reader images the multiplexed sensing membrane using the standard Android camera app (ISO: 50, shutter at 1/125, autofocused), and saves a raw image of the VFA sensing membrane (.dng file) for subsequent processing and quantification of the CRP concentration.

#### Data processing

Custom image processing software was developed to automatically detect and segment the immunoreaction spots in each mobile-phone image of the activated VFA cassette (see Fig. S2). After segmentation, the pixel average of each spot is calculated and subtracted by the pixel-average of a locally defined background containing BSA blocked NC membrane. Each background-subtracted spot signal is then normalized to the sum of all the spots on the sensing membrane. The final spot signal *s*′_*m,p*_ is therefore described by,

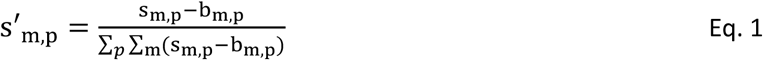

where *m* represents the spotting condition, and the *p* represents the *p*^th^ redundancy on the VFA per condition. *s_m,p_* is the pixel average of a given segmented spot, and *b_m,p_* is the local background signal. The final VFA signal per condition can then be calculated as:

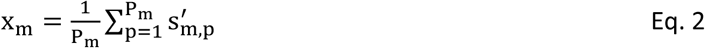

where *P_m_* is the number of redundancies for a given spotting condition. The normalization step in Eq. (1) helps us to account for sensor-to-sensor variations borne out of pipetting errors, fabrication tolerances, as well as operational variances.

### Clinical testing

We procured remnant human serum samples (under UCLA IRB #19-000172) for hsCRP testing using our VFA platform. Each clinical sample was previously measured within the standard clinical workflow as part of the UCLA Health System using the CardioPhase *hs*CRP Flex^®^ reagent cartridge (Cat. No. K7046, Siemens) and Dimension Vista System (Siemens). In total, we measured 85 clinical samples in triplicate with our VFA sensors. All but one sample was within the standard hsCRP range of 0 to 10 mg/L, with the outlier having a concentration of 83.6 mg/L. In addition to testing these clinical samples, nine CRP-free serum samples (Fitzgerald Industries International, 90R-100) were measured as well as nine artificial samples created by spiking 200, 500, and 1000 mg/L CRP into CRP-free serum samples. These artificial samples were tested to simulate serum samples from patients undergoing acute inflammatory events. Though relatively rare in the context of hsCRP testing, such high concentration samples can be falsely reported as having a low CRP concentration due to the hook-effect. Therefore, these samples were included to test if our multiplexed computational VFA could avoid such false reporting. Among different batches of 273 fabricated VFA sensors, we removed one VFA test from the data-set due to a fabrication error (misalignment, see Fig. S3a), and removed two triplicates due to abnormally high levels of nonspecific binding, which was immediately obvious in the low signals on the sensing membrane and unusual pink color observed on the top case (Fig. S3b).

### Computational VFA sensor analysis

After the clinical study was completed the image data from the activated VFA tests were partitioned into a training set (N_train_ = 209) and testing set (N_test_ = 57). This data partition was structured to ensure that the testing samples would be distributed linearly over the hsCRP range, and that samples were pulled proportionally from the different fabrication batches within each cardiovascular risk stratification group. The raw background-subtracted pixel average values are shown in Figure S4, where the marker color and shape indicate the fabrication batch ID and the reagent batch ID, respectively.

#### Model and cost function selection

The training set was analyzed via a k-fold cross-validation (k=5) to determine the optimal learning algorithm for quantification of CRP concentration from the inputs *X_IN_*. We evaluated different fully connected networks through a random hyper-parameter search, where the number of nodes, layers, regularization, dropout, batch-size, and cost-function were each randomly selected from a user-constrained list. A tiered neural network architecture (Fig. S5) with a cost function of mean-squared logarithmic error (MSLE) yielded the best performance over the random iterations of the crossvalidation. As an alternative, a single neural network with multiple hidden layers, in contrast to the tiered structure, could also be used in providing an accurate and generalizable model.

## RESULTS

### Optimization of VFA spots and conditions using machine learning

Machine learning-based optimization and feature selection of our VFA platform was performed in two distinct steps: spatial spot selection and condition selection, illustrated in Figure 2a and Figure 2b, respectively. For the spot selection process, a cost function, *j_m,p_*, was defined per sensing spot to represent the normalized distance from the mean of like-spots (*i.e*. spots that share the same condition) averaged over the samples in the training set,

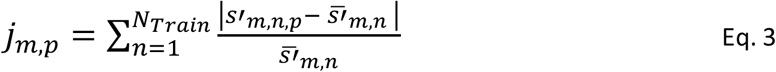

where *s’_m,n,p_* is defined in Eq. 1 with the added index *n* indicating the *n*^th^ sample in the training set. 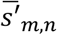 is the spot signal averaged over each condition within a single test, *i.e*. 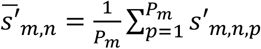.

**Figure 2.**
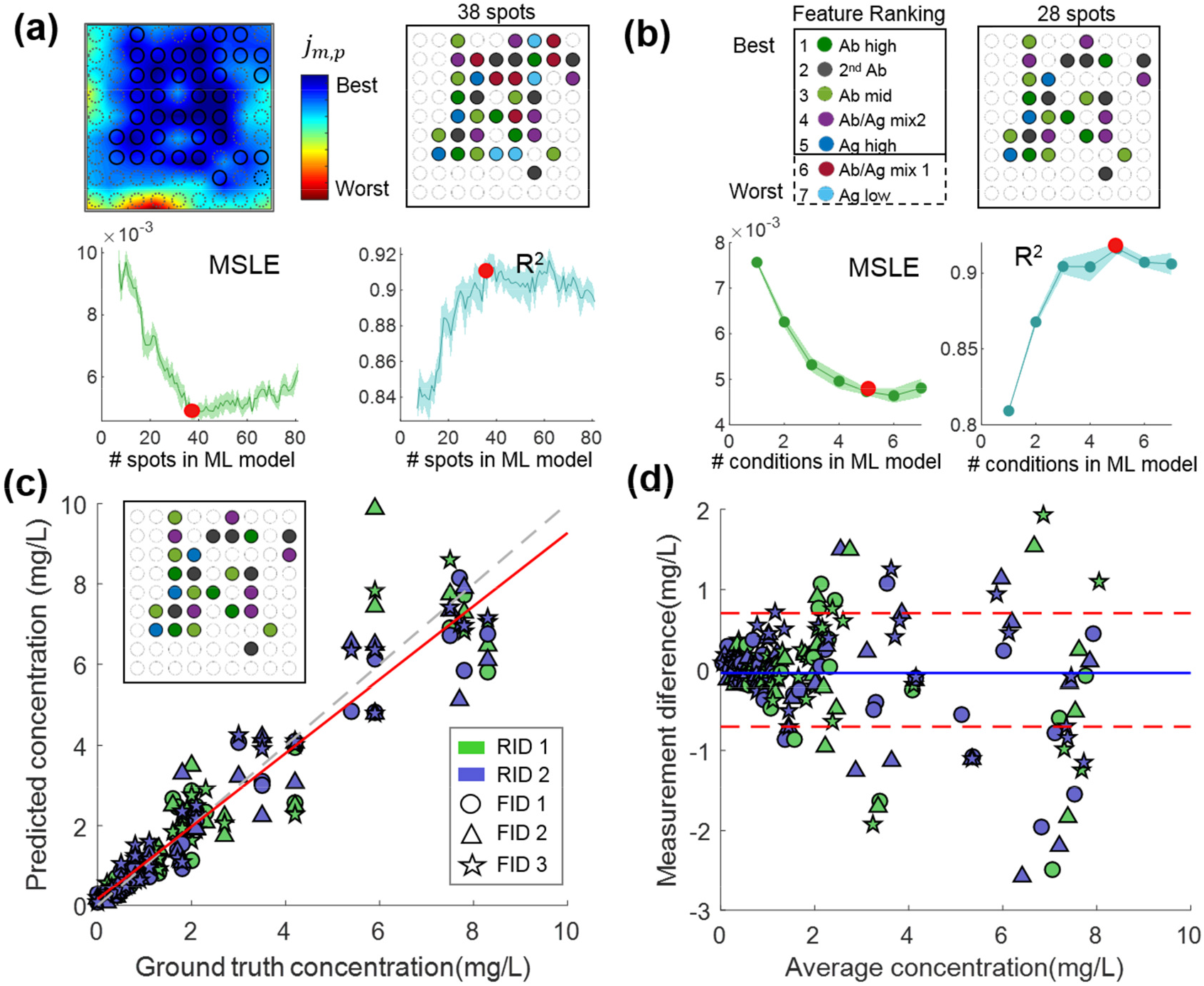
Cross-validation and feature selection analysis using the training data set of clinical samples (N_train_ = 209) (a) The spot selection process. A heat-map (top left) is generated by plotting the cost function *j_m,p_* across the sensing membrane. The cross-validation performance, both MSLE and the coefficient of variation (R^2^), is then plotted against the number of spots selected based off of *j_m,p_* (bottom). The optimal subset of spots (top right) is then selected based off the optimal quantification performance indicated by the solid red marker. (b) The condition selection process. Conditions are ranked based off of an iterative elimination method (top left), and the cross-validation performance is plotted against the number of conditions input into the quantification network. The optimal subset of conditions (top right) is then selected based off the optimal quantification performance indicated by the solid red marker. (c) The cross validation results using the selected features, where the ground truth CRP concentration is plotted against the predicted CRP concentration. The marker color and shape represent the different reagent batch ID (RID) and the fabrication batch ID (FID), respectively. (d) Bland-Altman plot of the same cross-validation results, where the dashed red lines represent the ± standard deviation of the measurement difference from the tested VFAs.

The heat map in Figure 2a, which is interpolated from a 9×9 matrix of the cost function defined at each spot of the VFA, visualizes the statistically robust active areas of the VFA sensing membrane. To select a subset of spots from the 9×9 grid configuration, we then performed a k-fold (k=5) cross-validation. The cross validation was performed over 75 iterations where the input to the neural network, *X_IN_*, was defined by incrementally smaller subsets of the original 81 spots for each iteration. The spot with the maximum cost *j_m,p_* was eliminated at each iteration, resulting in the last iteration containing a subset of 7 spots, each corresponding to a different condition. The MSLE value from the cross validation was then plotted for every iteration to visualize the trade-off between the number of spots and the error of the network inference (Fig. 2a). Due to the random training process of the neural network, there is noise associated with this curve, however a clear performance benefit can be seen after the elimination of the first 30 to 40 spots corresponding to the highest *j_m,p_*. It is also clear that further reducing the number of spots results in substantial increase in quantification error. Therefore, the approximate minimum of the MSLE curve was used to define a subset of 38 spots for subsequent analysis.

After this initial spot selection (Fig. 2a), this subset of 38 spots was further subject to a condition selection step to further optimize the performance of our computational VFA for hsCRP. This second phase of the feature selection aims to select the most robust sensing channels as defined by the unique chemistry attributed to the different spotting conditions. To this end, we performed a second iterative k-fold (k=5) cross-validation analysis, eliminating one spotting condition each iteration and tracking the cross-validation error as a result of each elimination. This process was repeated for incrementally smaller subsets of conditions defined by the minimum MSLE result from the previous iteration. Resulting from this analysis, Fig. 2b reports the MSLE and coefficient of determination as function of the number of spotting conditions, suggesting that eliminating the Mix 1 and Ag-low condition can lead to slightly better or equivalent performance when compared to the inclusion of all the original spotting conditions.

Taken together, this machine learning-based optimization of the VFA leads to the statistical selection of the best combination of spots and conditions (Fig 2c inset) that can computationally determine the analyte concentration. The cross-validation results, compared to the gold standard hsCRP measurements, are also reported in Figs. 2c and d. Here the inputs to the neural network, *X_IN_*, are defined by the optimal spot configuration as determined by the spot and condition selection (see Fig 2c inset), and also include two additional integer features which correspond to the reagent ID (*RID* ∈ {0,1}) and the fabrication batch ID, (*FID* ∈ {1,2,3}).

After this feature selection and cross-validation analysis reported in Figure 2, the final CRP quantification algorithm was trained using the entire training set (N_train_=209) and the optimal spot configuration (Fig 2c inset). In addition to the CRP quantification algorithm, a second classification algorithm was trained to identify the CRP samples representing an acute inflammation event, with a CRP concentration of > 10 mg/L 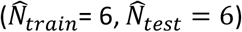 (Fig 1d). Next, we report our blind testing results using this optimized CRP VFA platform.

### Validation of computational VFA performance for CRP measurements

Our computational VFA results from the blind testing set (N_test_=57) correlated well to the quantification results of the gold-standard hsCRP Flex cartridge run on the Dimension Vista System (see Figure 3).

**Figure 3.**
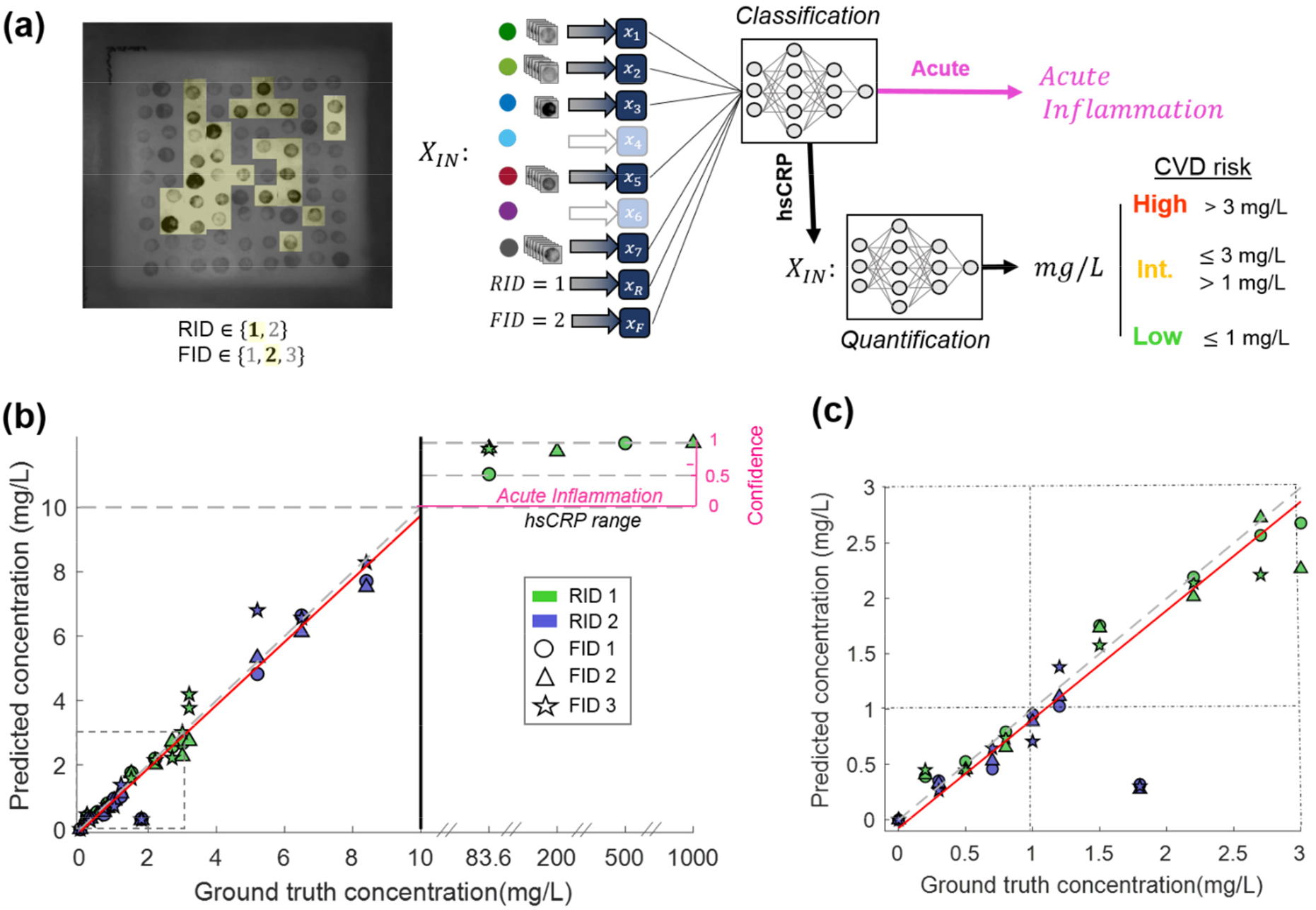
Blind testing results of clinical samples (N_test_ = 57) (a) The features selected from the crossvalidation analysis are extracted from a blind testing image and input into the neural network-based processing which infers the final CRP concentration. The clinical cutoffs for stratifying patients in terms of cardiovascular disease (CVD) risk are shown on the right. (b) The ground truth CRP concentration plotted against the VFA predicted CRP concentration (left y-axis) from blindly tested clinical samples. The dotted line represents a perfect match (y = x) and the red line represents the linear best fit. The confidence score is plotted (right y-axis) for the samples classified as acute. The marker color and shape represent the different reagent batch ID (RID) and fabrication batch ID (FID), respectively. (c) The blind testing results for the low and intermediate CVD risk regimes, where the dotted lines represent the clinical cutoffs at 1 and 3 mg/L.

These samples were analyzed using only the pixel information contained within the computationally determined subset of 28 spots and 5 conditions (Fig. 3a). The *x_m_* signals (Eq. 2) along with the FID and RID of each test sample were first classified by an initial neural network to determine if the test was in the hsCRP range (<10 mg/L) or the acute inflammation range (> 10 mg/L); we achieved 100% classification accuracy, and correctly classified 6 samples as acute and the rest (51 samples) as in the hsCRP range. The samples classified in the hsCRP range were then routed to a quantification neural network, whereas the acute samples were simply reported as acute along with a confidence score, as summarized in Fig. 3c.

The quantification accuracy of the hsCRP samples using our computational VFA was characterized by a direct comparison to the gold-standard values (Figs. 3b-c). With 51 tests quantified in the hsCRP range, the R^2^ value was found to be 0.95, with a slope and intercept of the linear best-fit line being 0.98 and 0.074 respectively. The overall average CV of the blind testing data was found to be 11.2% with the average CV for the low-risk, intermediate-risk, and high-risk stratified samples quantified as 11.5%, 10.1%, and 12.2 %, respectively. As a reference point, the FDA review criteria for hsCRP testing state an acceptance criterion of ≤ 20% overall CV, with a specific CV of ≤ 10% for samples in the low risk category (*i.e*. < 1 mg/L) ^38^.

## DISCUSSION

Our VFA-based hsCRP test benefits from machine learning in several ways. Firstly, using neural networks to select optimal spots and infer analyte concentration from the highly multiplexed sensing channels greatly improves our quantification accuracy when compared to *e.g*. a standard multi-variable regression (see Fig. S6). Deep learning algorithms such as the fully-connected network architecture used in this work, contain a much larger number of learned/trained coefficients along with multiple layers of linear operations and non-linear activation functions when compared to standard linear regression models. These added degrees of freedom enable neural networks to converge to robust models which can learn non-obvious patterns from a confounding set of variables, making them a powerful computational tool for assay interpretation and calibration. However, one concern with deep learning approaches is the possibility of overfitting to the given training set, especially in the instance of limited data. To mitigate this issue, we incorporated regularization terms in the hyper-parameter search (both L2 regularization and dropout), and found via cross-validation that the lowest error model employed the maximum degree of dropout regularization (*i.e*. 50%) ^46,47^. However, we observed better quantification results in the blindly tested samples when compared to the cross validation analysis, suggesting that our model appropriately generalized over the operational range of the hsCRP sensor.

Secondly, by incorporating fabrication information using *RID* and *FID* input features, the neural network was able to learn from batch-specific patterns and signals. This resulted in a 12.9% reduction in the blindly tested MSLE when compared to the performance of a network trained without these fabrication batch input features. Similarly, incorporating the fabrication information reduced the overall CV from 16.64% to 11.2% and increased R^2^ value from 0.92 to 0.95. It is important to note that these VFA tests (N=273) were fabricated without the use of industry-grade production equipment such as humidity and temperature controlled chambers, and in addition, several fabrication steps involved manual assembly. Taken together, these simple input features can benefit the performance and quality assurance of future computational POC tests following the methodology of this work. For example, the fabrication information could be included for each test in the form of a Quick Response (QR) code or could alternatively be logged into a GUI by the user before the measurement data are sent to the quantification network (running on a local or remote computer).

Another benefit of our computational VFA platform is the mitigation of false sensor response due to the hook effect. The VFA format importantly enables rapid computational analysis of highly multiplexed immunoreaction spots with minimal cross talk or interference among spots, which is inevitable for the case of standard lateral flow assays or RDTs. The multiplexed information reported by the different spotting conditions therefore allows for unique combinatorial signals to be generated over a large dynamic range (see Fig. 3b). The hook effect is clearly seen in our raw sensor data, exhibited by the capture antibody (Ab) condition (see Fig S1, Fig. S4), illustrating how this condition alone can lead to false reporting of high analyte concentrations, *i.e*. in the case of acute inflammation. Therefore, without the incorporation of the monotonically responsive CRP antigen (Ag) spotting condition as one of the multiplexed channels in the VFA, high-concentration CRP samples can be falsely reported as low concentration due to the hook effect. This conclusion would still be true even if we trained another neural network that used a limited number of conditions as input; for example, by re-training the classification network using only the Ab and Secondary Ab spotting conditions as inputs, we found that the 83.6, 200, and 1000 mg/L samples are falsely reported as having CRP concentrations of 7.81, 7.34, and 3.84 mg/L respectively. In the case of analyzing only the Ab channel, all of the high-concentration CRP samples would have been falsely reported as having concentrations below 10 mg/L. These results highlight the importance of multiplexed sensing in our computational VFA platform to mitigate the limitations induced by the hook effect in order to algorithmically enhance the dynamic range of our sensor.

### Computational sensing for assay development

Computational sensing broadly refers to the joint design and optimization of sensing hardware and software, and as implemented in this study, provides a framework for data-driven assay development where the diagnostic or quantification algorithm informs the multiplexed sensor design and vice versa. As detailed in the Methods section, the computational sensing approach begins with the selection of a neural network architecture and associated cost function. This first step is paramount to the computational sensor design, as it defines the model and error metric with which the subsequent feature selection is performed. The determination of the cost function therefore poses an interesting question for future computational sensors and diagnostic tests: because the selection of the cost function defines the training of a neural network, what are the most clinically appropriate error functions with which one should design a computational sensing system? For example, in the case of cardiovascular risk stratification with the hsCRP test, an error of ± 0.1 mg/L is more problematic for samples that are in the range of the clinically defined cutoffs (*i.e*. 1 and 3 mg/L) when compared to samples with relatively higher CRP concentrations, such as 8 mg/L. Therefore, a traditional cost function for regression such as the mean-squared-error may not be as appropriate as the mean-squared-logarithmic-error or mean-absolute-percentage error, which take into account the relative ground-truth concentration for each error calculation. Therefore, special consideration must be given to the cost functions employed, and custom cost functions defined jointly by physicians/clinicians and engineers should be considered.

Feature selection and machine learning based optimization can similarly be used to inform the sensing membrane design. POC sensors can especially benefit from feature selection to circumvent noise borne out of their low-cost materials (such as paper used in our VFA) and operational variations. For example, the heat-map in Fig. 2a very well reveals how the immunoreaction spots closest to the edges of the sensing membrane contain the most variation in their normalized signals. This most likely results from the position-dependent vertical flow variations inherent in the inexpensive VFA format, which uses paper materials totaling <$0.2 per CRP test (Table. S1). These areas can therefore be avoided in future iterations of the sensor development, saving reagent costs and fabrication time, while also preserving robust sensing channels. Furthermore, identifying these areas of statistical variation can also inform the fabrication process. For example, Fig. 2a also shows that the top edge of the VFA sensing membrane as statistically more robust than the bottom and sides of the sensing membrane. Therefore, this spot selection analysis indicates a unidirectional fabrication bias in the lateral alignment of the sensing membrane within the VFA stack, which can be addressed in future iterations of the batch fabrication process.

Complementing the spot selection, the statistical condition selection process investigates the efficacy of the sensing channels and the unique immunoreactions defined by their spotting condition. Inherent complexities of the underlying chemistry such as the stochastic arrangement of the capture proteins within the porous NC membrane, as well as the effects of steric hindrance, pH, humidity, and temperature can obscure intuition behind the selection of spotting conditions for a given sensing application. Therefore, computational sensing systems can benefit from data-driven selection of sensing channels. For example, Fig. 2b shows that the quantification performance improves slightly upon the out-right elimination of the Mix 1 and Ag-low conditions. This suggests that their signal response is redundant or less stable when compared to the other conditions, and is confirmed by the poor repeatability of the Ag signal between the reagent and fabrication batches (see Fig. S4). Such a feature selection procedure in a highly multiplexed format like the VFA could therefore be used to computationally screen spotting conditions from a large number of differing capture chemistries including, but not limited to, different structures of capture antibodies/antigens (i.e., polyclonal vs. monoclonal) as well as varying buffer conditions and reagent concentrations. Conditions which do not empirically benefit sensor performance can be replaced by new conditions in another iteration of the development phase, or be replaced by additional redundancies of effective conditions in order to benefit from signal averaging.

Additionally, this statistical feature selection and optimization process can inform cost-performance trade-offs to help design the most robust and cost-effective implementations of POC assays. For example, the reagent cost for the immunoreaction spots contained in the hsCRP VFA test is reduced by 62%, from $2.61 to $0.97 per test, by implementing only the computationally selected chemistries. Additionally, certain spotting conditions might have an optimal capture protein concentration due to steric hindrance effects or higher degrees of nonspecific binding. Therefore, in a computational sensor, reagent costs can be significantly reduced without sacrificing assay performance by employing these statistically optimized capture-protein concentrations. One should also note here that these reagent costs per test would be significantly reduced under large scale manufacturing, benefiting from economies of scale, which is expected to bring the total cost per test (including all the materials and reagents) to <$0.5.

Taken together, we showed a data-driven sensor design and read-out framework, enabled by deep learning, for improving POC tests. As a use-case scenario, we demonstrated hsCRP testing with a colorimetric paper-based multiplexed VFA and clinical samples covering a large dynamic range. The multiplexed sensing membrane contained in the VFA was jointly developed with a quantification algorithm based on a fully-connected neural network architecture. First, a training data-set was formed by measuring human serum samples with the VFA. Then, through cross-validation of the training set, the most robust subset of sensing channels was selected from the multiplexed sensing membrane and used to train a CRP quantification network. The network was then blindly tested with additional clinical samples and compared to the gold standard CRP measurements, showing very good agreement in terms of quantification accuracy and precision. Additionally, the multiplexed channels and computational analysis helped us overcome limitations to the operational range of the CRP test borne out of the hook-effect. Our results demonstrate how a computational sensing framework and multiplexed sensor design can be used to engineer robust and cost-effective POC tests that have the potential to democratize diagnostics and expand access to care.

## Acknowledgments

The authors would like to acknowledge the NSF PATHS-UP Engineering Research Center and HHMI for funding. The authors would also like to acknowledge the Molecular Screening and Shared Resource at the California NanoSystems Institute (CNSI), and Dr. Robert Damoiseuax of UCLA for the assistance with the protein spotting.

## Contributions

Zachary Ballard optimized and fabricated VFA sensing platform, performed the clinical testing measurements, and developed the computational sensing framework and data analysis. Hyou-Arm Joung optimized and fabricated the VFA sensing platform, performed the clinical testing measurements including the reagent handling and synthesis. Artem Goncharov wrote the automated image processing code for sensor analysis. Jesse Liang worked to optimize and develop the protein spotting process. Karina Nugroho optimized and fabricated the VFA sensors. Omai Garner, Dino Di Carlo, and Aydogan Ozcan oversaw and supervised the research. Aydogan Ozcan initiated and conceptualized the computational sensing project.

